# Neural Deterioration and Compensation in Visual Short-term Memory Among Individuals with Amnestic Mild Cognitive Impairment

**DOI:** 10.1101/2023.11.19.567711

**Authors:** Ye Xie, Wei Zhang, Yunxia Li, Yixuan Ku

**Affiliations:** Guangdong Provincial Key Laboratory of Brain Function and Disease, Center for Brain and Mental Well-Being, Department of Psychology, Sun Yat-sen University, Guangzhou, China; Department of Neurology, Tongji Hospital, School of Medicine, Tongji University, Shanghai, China; Department of Neurology, Shanghai Pudong Hospital, Fudan University Pudong Medical Center, 2800 Gongwei Road, Pudongm, Shanghai 201399, China; Shanghai Key Laboratory of Vascular Lesions Regulation and Remodeling; Peng Cheng Laboratory, Shenzhen, China

## Abstract

Amnestic mild cognitive impairment (aMCI) is considered to carry a high risk of progression to Alzheimer’s Disease (AD) and it has been characterized by deficits in visual short-term memory (VSTM). However, the relationship between VSTM deficits and pathological brain damage in individuals with MCI has remained unknown. In the current study, we examined a group of 123 elder adults, including 55 with aMCI and 68 age-matched controls. Participants performed color change-detection VSTM tasks, and structural and functional magnetic resonance images were acquired during rest. Compared to the normal control (NC) group, individuals with aMCI exhibited poorer accuracy and longer reaction time in VSTM tasks, along with reduced VSTM capacity. Additionally, structural atrophy was observed in aMCI participants in several brain regions, including the left medial temporal lobe (MTL), the left thalamus, the right frontal pole (FP) and the right postcentral gyrus (postCG). Interestingly, VSTM accuracy and capacity were found to be associated with the volume of the left MTL in the NC group but not in the aMCI group, suggesting alterations in the relationship between VSTM and brain regions in aMCI. However, VSTM capacity was correlated with the volume of the right FP in both groups, suggesting potential compensatory mechanisms involving the prefrontal cortex in aMCI. Moreover, using the atrophic left MTL as a seed, functional connectivity to the right FP was significantly higher in aMCI compared to NC. Notably, this FP area showed overlap with the atrophic frontal areas in terms of structural abnormalities. Furthermore, for individuals with aMCI who had a larger left MTL, the compensatory involvement of the right FP in VSTM, as assessed by brain-behavior correlations, was diminished. In summary, the present study uncovered a mechanism involving MTL dysfunction and prefrontal compensation in aMCI when performing VSTM tasks. These findings may offer valuable insights into potential intervention targets for individuals with aMCI.

## Introduction

Alzheimer’s Disease (AD) is an irreversible neurodegenerative condition that significantly impacts daily life of elderly individuals and imposes substantial societal burdens ^1-5^. The hallmark characteristics of this disease include brain atrophy and cognitive impairment. Amnestic mild cognitive impairment (aMCI) represents a transitional stage in the progression to AD, with neuropathological features that lie between the neuroplasticity observed in normal aging and the pathological changes seen in AD ^6,7^. Therefore, investigating behavioral markers and neuropathology during this period is of utmost importance for early diagnosis and intervention in AD^1,8^.

Recent research has identified deficits in short-term memory, especially in visual short-term memory (VSTM), as a sensitive behavioral indicator of the early stages of AD. These VSTM deficits have been observed in AD and its early stages, as well as in asymptomatic carriers of familial AD, especially when feature binding is engaged ^9-12^. Notably, VSTM binding performance has been shown to distinguish early MCI from healthy controls ^13^. Moreover, a recent study reported an association between VSTM performance and tau and ß-amyloid burdens in familial AD carriers (MCI and asymptomatic carriers) ^14^. Given these findings suggesting VSTM as an early marker of AD pathology, a crucial question arises: how is VSTM influenced by brain deterioration during the aMCI stage?

Previous literature has established that VSTM relies on several brain regions, including the prefrontal cortex ^15^. The prefrontal cortex is implicated in providing top-down control, maintaining task-relevant information, and filtering out irrelevant distractions during VSTM processes ^16,17^. Prefrontal engagement has been closely linked to VSTM capacity ^18^. As cortical atrophy is believed to progress from limbic and temporal cortices to higher-order association areas and eventually primary sensorimotor regions in AD ^19^, the neural networks supporting VSTM may ultimately be affected, leading to VSTM deficits.

Hippocampus, a critical brain region for memory function and one of the most vulnerable regions to AD pathology ^20^, is of particular interest when addressing this question. While early reports suggested that damage to the medial temporal lobe (MTL), including the hippocampus, primarily affected long-term memory rather than short-term memory ^21^, recent studies have increasingly implicated the important role of hippocampus in VSTM ^16,17,22^. Recent research has identified specific neurons with persistent firing during the maintenance phase of VSTM tasks, with their firing patterns associated with task workload ^23^. Another study found that maintaining VSTM representations of task was tied to the phases of low-frequency activity in hippocampus ^24^. These studies highlighted the critical role of the hippocampus in supporting VSTM, raising the possibility that early deterioration in MTL regions might influence VSTM performance in aMCI individuals.

To investigate the relationship between neural deterioration and VSTM deficit in aMCI individuals, we enrolled 55 participants with aMCI and 68 age-matched normal elder controls (NC) in the current study (see Table 1 for demographic characteristics). We conducted structural and resting-state functional magnetic resonance imaging (MRI) assessments and evaluated their VSTM using a change-detection task (Fig. 1) ^25-27^. Firstly, we would examine VSTM deficits in aMCI. Secondly, we would investigate the relationship between VSTM performance and brain regions displaying atrophy in aMCI. We hypothesize that the support provided by the MTL to VSTM might be impaired in the aMCI group and the neural networks related to VSTM would still be capable of supporting VSTM, dependent on the severity of disease. By addressing these objectives, we aim to shed light on the relationship between neural deterioration and VSTM deficits in aMCI, contributing to a better understanding of the early stages of AD and potentially informing intervention strategies.

**Figure 1.**
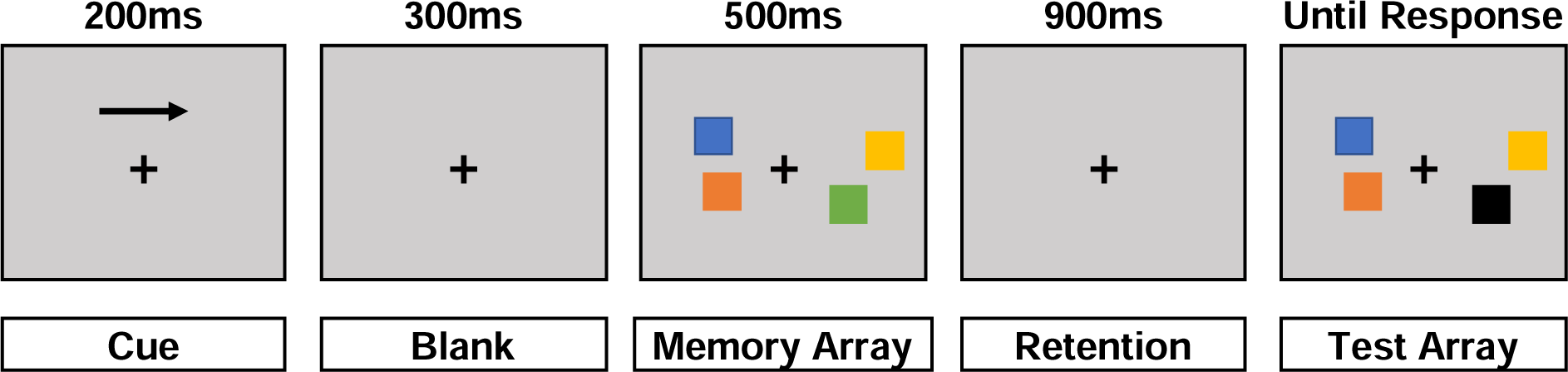
Stimuli and trail procedure of visual change detection task.

**Table 1.** demographic characteristics.

|  | aMCI<br>(n = 55) | NC<br>(n = 68) | Between-group differences |
| --- | --- | --- | --- |
| Age (years) | 72.42 (6.69) | 70.31 (6.19) | $t(121) = 1.81, p = 0.072^a$ |
| Gender (male: female) | 23:32 | 44:24 | $\chi^2 = 6.42, p = 0.011^b$ |
| Education (years) | 11.33 (2.65) | 12.43 (2.79) | $t(121) = -2.22, p = 0.028^a$ |
| MMSE | 24.86 (2.48) | 27.47 (1.65) | $W = 638.5, p < 0.001^c$ |
| MoCA | 17.07 (3.34) | 24.24 (2.48) | $t(121) = -13.64, p < 0.001^a$ |

Values are mean (standard deviation). Abbreviation: aMCI, amnesic mild cognitive impairment; NC, normal controls; MMSE, Mini-Mental State Examination; MoCA, Montreal Cognitive Assessment.
<sup>a</sup> independent two-sample t-test, <sup>b</sup> chi-square test, <sup>c</sup> Mann-Whitney U test.

## Result

### VSTM Performance

The 2 (group: aMCI or NC) x 2 (load: 2-item or 4-item) repeated-measures ANOVA was applied on the ACC and RT of change detection task with age, gender, education and collecting site as covariates.

For ACC, the ANOVA test revealed a significant main effect of group (F(1,117) = 15.38, *p* < 0.001; Fig. 2A). *Post hoc t* tests using Bonferroni correction showed significantly lower ACC for the aMCI group than the NC group in both 2-item (*p* < 0.001) and 4-item (*p* = 0.014) change detection task.

For RT, the ANOVA test revealed a significant main effect of group (F(1,117) = 5.26, *p* = 0.024; Fig. 3B) and a significant interaction between group and load (F(1,117) = 8.62, *p* = 0.004; Fig. 3B). *Post hoc t* tests using Bonferroni correction showed significantly longer RT for the aMCI group than the NC group in 2-item (*p* = 0.002) change detection task.

Capacity was calculated based on the performance of 4-item change detection task. ANCOVA test was used to compare capacity with age, gender, education and collecting site as covariates. The results showed significant lower capacity for the aMCI group than the NC group (*p* = (F(1,117) = 8.382, *p* = 0.005; Fig. 2C).

**Figure 2.**
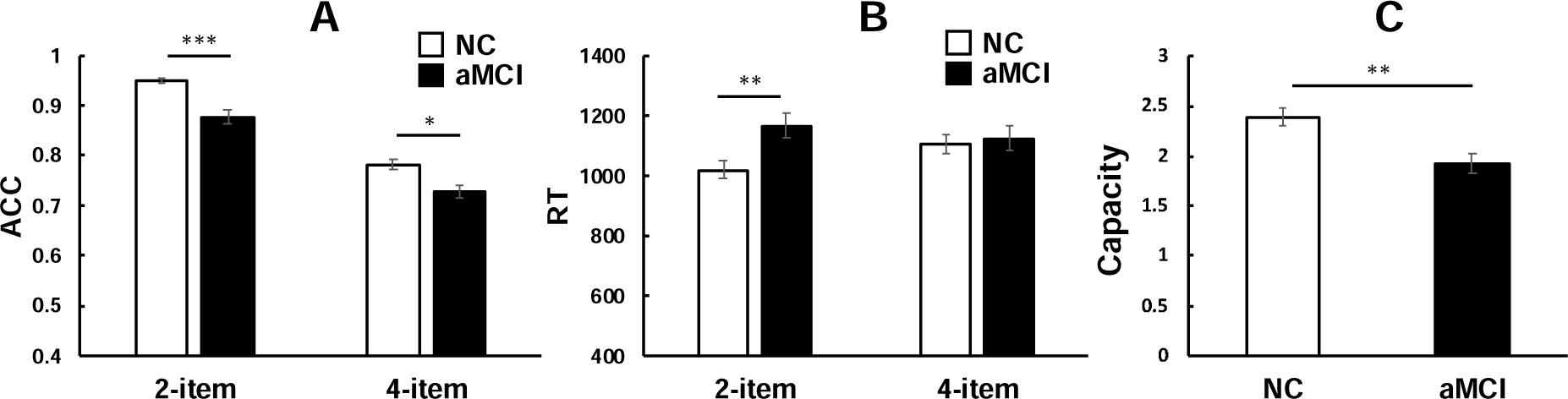
performance of change detection task. Error bar represented standard error. ACC, accuracy; RT, reaction time; aMCI, amnestic mild cognitive impairment; NC, normal control. *, *p* < 0.05; **, *p* < 0.005; ***, *p* < 0.001.

### Structural difference and correlation with VSTM

Two-sample t-test showed that the gray matter volume of left thalamus (peak: -18, -33, 6), right postcentral gyrus (postCG; peak: 44, -18, 38), left medial temporal lobe (MTL; including hippocampus/amygdala area; peak: -27, -10, -16) and right frontal pole (FP; peak: 24, 54, 14) were significant smaller in the aMCI group when comparing with the NC group (Fig. 3A and Table 2, voxel threshold *p* < 0.005, cluster threshold *p* < 0.05 by GRF correction, with TIV, age, gender, education and collecting site as covariates). We further extracted the gray matter volume from each of the clusters and all the clusters showed significantly smaller gray matter volume in the aMCI group compared to the NC group (*ps* < 0.005; Fig. 3B).

We further test the correlation between the gray matter volume of these brain regions and VSTM performance for each of the aMCI group and NC group. TIV, age, gender, education and collecting site were used as covariates. The results showed that the gray matter volume of left MTL was significantly positively correlated with ACC of both 2-item and 4-item change detection task and capacity of 4-item change detection task in the NC group but no significant correlation was found in the aMCI group (Fig. 3C). The gray matter volume of right FP was significantly positively correlated with ACC and capacity of the 4-item change detection task in both the aMCI group and the NC group (Fig. 3D). No significant correlation was found between VSTM performance and gray matter volume of left thalamus or right postCG (*ps* > 0.05).

**Figure 3.**
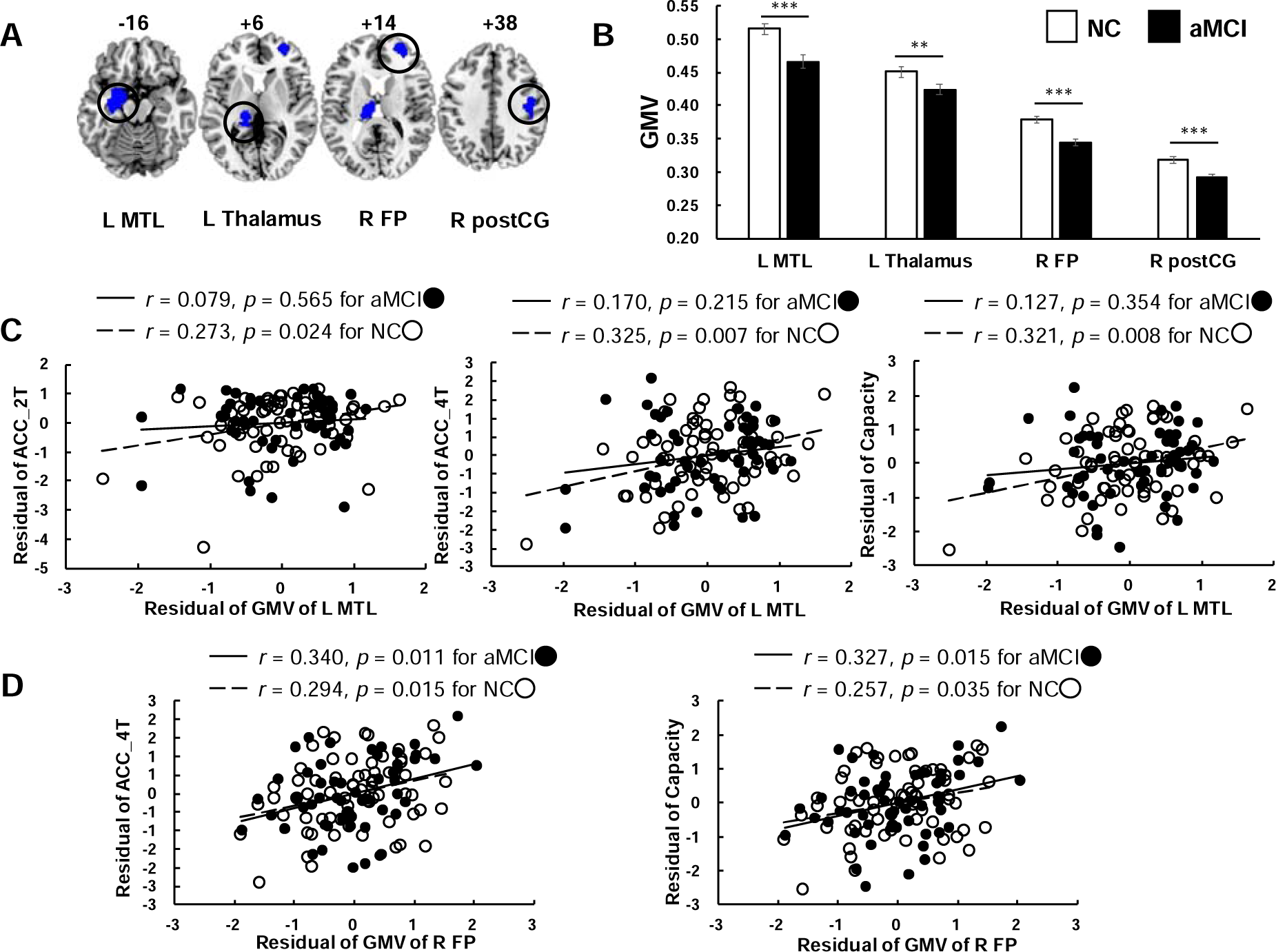
(A) structural atrophy in aMCI group compared to NC group. Voxel threshold *p* < 0.005, GRF correction with cluster threshold *p* < 0.05, gray matter mask applied and age, gender, education, TIV and collecting site as covariates. Error bar represented standard error. **, *p* < 0.005; ***, *p* < 0.001. (B) Significant Structural differences between NC and aMCI group. (C) Relationship between structural gray matter volume (GMV) of left MTL and performance of change detection task for each group. Age, gender, education and collecting site were used as covariates. (D) Relationship between structural GMV of right FP and performance of change detection task for each group. Age, gender, education and collecting site were used as covariates. Error bar represented standard error. Abbreviation: aMCI, amnestic mild cognitive impairment; NC, normal control; FP, frontal pole; HP, hippocampus; MTL, medial temporal lobe; postCG, postcentral gyrus; L, left; R, right.

**Table 2.** Brain regions showed significant structural differences between aMCI group and NC group.

| Region | Cluster Size<br>(voxels) | Peak t-score | Peak MNI Coordinate<br>mm (x, y, z) |
| --- | --- | --- | --- |
| ***aMCI < NC*** |  |  |  |
| L Thalamus | 1269 | 4.95 | -18, -33, 6 |
| R postCG | 1602 | 4.51 | 44, -18, 38 |
| L HP/AMG | 2928 | 4.37 | -27, -10, -16 |
| R FP | 1389 | 4.24 | 24, 54, 14 |
| ***aMCI > NC*** |  |  |  |
| No brain region |  |  |  |

### Functional connectivity difference and correlation with VSTM

We further used the left MTL cluster and the right FP cluster, which gray matter volume was correlated to the VSTM performance, as ROI seed to test whether the functional circuits of MTL or prefrontal regions would be influenced by disease and how’s their association with VSTM in aMCI participants (Fig. 5A). The seed-to-whole brain functional connectivity analysis revealed right FP (peak: 16, 46, 26) as target regions which showed significant greater functional connectivity with left MTL cluster in aMCI group when compared to NC group (Fig. 5B and Table 3, voxel threshold *p* < 0.005, cluster threshold *p* < 0.05 by GRF correction, with age, gender, education and collecting site as covariates). This brain area was adjacent to the right FP area which showed significant atrophy in aMCI group (Fig. 5C: red area represented the current area identified in functional analysis, blue area represented the area identified in structural analysis, green area represented the overlap of these two areas). The *β* value was extracted from the current right FP and the independent-sample t-test showed significant greater functional connectivity with left MTL in the aMCI group than the NC group (Fig. 5D). No significant correlation was found between the functional connectivity circuits of left MTL and performance of change detection task in either group. No significant cluster was found in the functional connectivity comparison analysis for right FP cluster identified in the structural analysis as ROI seed (Table 3, voxel threshold *p* < 0.005, cluster threshold *p* < 0.05 by GRF correction, with age, gender, education and collecting site as covariates).

**Figure 4.**
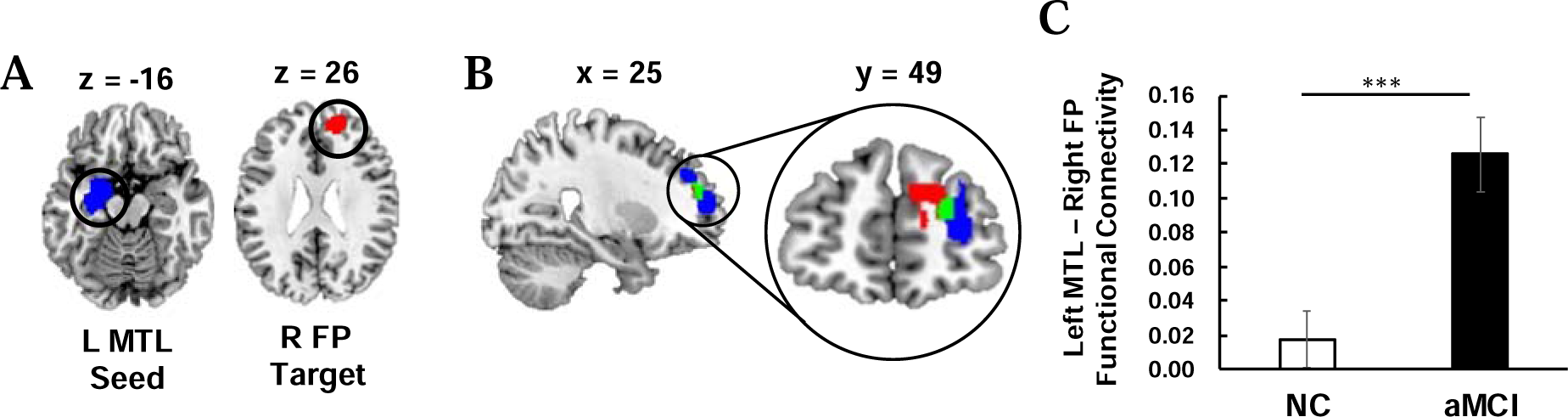
(A) R FP (target) area showed significant between-group difference in functional connectivity to L MTL (ROI seed). Voxel threshold *p* < 0.005, GRF correction with cluster threshold *p* < 0.05, gray matter mask applied and age, gender, education and collecting site as covariates. **, *p* < 0.005; ***, *p* < 0.001. (B) display of the right FP areas. Red, the right FP in functional results; blue, the right FP in structural results; green, overlap area. (C) Significant between-group difference in functional connectivity between left MTL and right FP. Error bar represented standard error. Abbreviation: aMCI, amnestic mild cognitive impairment; NC, normal control; FP, frontal pole; MTL, medial temporal lobe; L, left; R, right.

**Table 3.** brain regions showed significant functional connectivity difference to ROI seeds between aMCI group and NC group.

| Region | Cluster Size<br>(voxels) | Peak t-score | Peak MNI Coordinate<br>mm (x, y, z) |
| --- | --- | --- | --- |
| ***L MTL*** |  |  |  |
| ***aMCI < NC*** |  |  |  |
| No brain region |  |  |  |
| ***aMCI > NC*** |  |  |  |
| R FP | 570 | 4.17 | 16, 46, 26 |
| ***R FP*** |  |  |  |
| ***aMCI < NC*** |  |  |  |
| No brain region |  |  |  |
| ***aMCI > NC*** |  |  |  |
| No brain region |  |  |  |

### The interaction between MTL and frontal area

As the significant correlations between the VSTM performance and gray matter volume of right FP were found in both groups, we were interested in if the underlying mechanism of such correlation was different between the two groups. Thus, the influence of left MTL atrophy on the relationship between the VSTM performance and frontal area was examined for each group. Participants were split into two subgroups, larger gray matter volume of left MTL (LV subgroup) and smaller gray matter volume of left MTL (SV subgroup) by comparing with the median of gray matter volume of left MTL for each group. The correlation between VSTM performance and right FP was conducted in each subgroup. The results showed that in the aMCI group, significant correlation between VSTM performance and gray matter volume of right FP was found in the SV subgroup (Fig. 5), while in the NC group, significant correlation between VSTM performance and gray matter volume of right FP was found in the LV subgroup (Supplementary Fig. 1).

**Figure 5.**
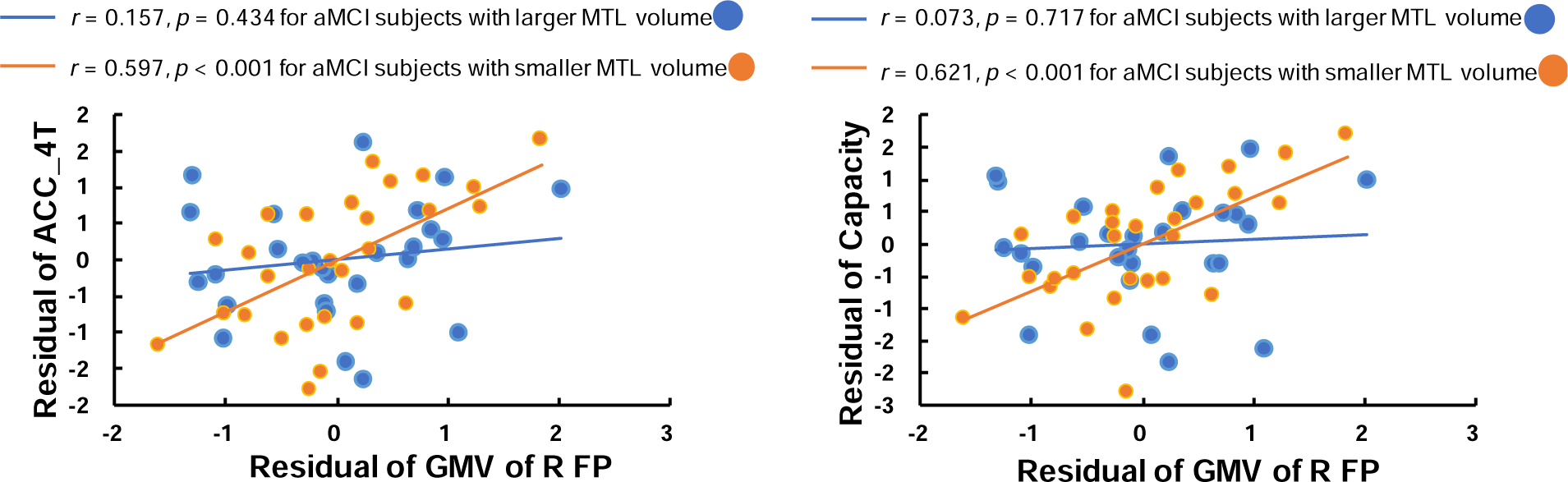
Relationship between gray matter volume (GMV) of right FP and VSTM performance influenced by the GMV of left MTL in the aMCI group. Abbreviation: aMCI, amnestic mild cognitive impairment; FP, frontal pole; MTL, medial temporal lobe; L, left; R, right.

As the right FP area targeted in the functional circuits of the left MTL was adjacent to the atrophic right FP area in the structure, we further test if the functional connectivity between these two areas would be associated with gray matter volume of left MTL. The correlation analysis revealed the functional connectivity between left MTL and right FP was marginally significant correlated with the gray matter volume of left MTL (*r* = 0.290, *p* = 0.063).

## Discussion

In the current study, we conducted a comprehensive examination of individuals with aMCI and age-matched normal elderly controls, shedding light on the intricate relationship between cognitive deficits, structural brain atrophy, and potential compensatory mechanisms. Specifically, people with aMCI performed worse in VSTM tasks compared to the NC group, together with structural atrophy in specific regions, including the left medial temporal lobe (MTL), right frontal pole, left thalamus, and right postcentral gyrus. Interestingly, in the NC group, VSTM performance correlated with the size of the medial temporal lobe. However, this correlation was absent in the aMCI group. Moreover, VSTM performance was linked to the size of the right frontal pole in both groups, and the right frontal pole played a more prominent compensatory role in VSTM tasks for aMCI individuals with smaller left MTL.

VSTM impairment has been consistently observed in individuals with early-stage AD, including asymptomatic carriers of familiar AD. This suggests the potential utility of VSTM as a cognitive marker for early AD detection ^10,28-30^. In the present study, the aMCI group exhibited poorer accuracy and slower reaction time, as well as reduced VSTM capacity, aligning with prior findings. These results underscore the presence of VSTM deficits in preclinical stages of AD.

Consistent with the earlier research, structural brain imaging in the aMCI group revealed atrophy in regions such as the MTL, thalamus and frontoparietal areas, reflecting the wide-spread neurodegeneration seen in the early stage of AD ^31^. Notably, the left MTL atrophy has consistently served as a robust neurostructural biomarker for predicting the progression from aMCI to full-blown AD ^32^. In the current study, the gray matter volume of the left MTL was correlated with VSTM performance in the NC group but not in the aMCI group. MTL, especially the hippocampus, has been proposed to play an important role in VSTM ^23,24,33,34^. Our results provide additional evidence supporting the role of MTL in VSTM and indicate dysfunction within the MTL during the early phases of AD, aligning with our initial hypothesis.

Regarding the neural networks involved in VSTM, we observed the frontal atrophy in the aMCI group, which aligns with prior literature reporting frontal atrophy in individuals with MCI ^3,7^. It has been proposed that the brain atrophy progresses from limbic and temporal areas to higher-order association regions such as the prefrontal cortex during the course of AD ^19^. Moreover, aMCI individuals with greater loss in the MTL and frontal areas are more likely to progress to AD ^20^. In addition, we found a positive correlation between gray matter volume in the right FP and the accuracy and capacity of VSTM in both groups. The prefrontal cortex is known to play a role in the VSTM process ^16,17,35^, which may explain this correlation observed in both groups. However, when we furtherly investigated this correlation by dividing participants into subgroups based on the volume of left MTL, we observed different mechanisms in each subgroup. The coherence between the prefrontal area and MTL has been regarded as an important mechanism supporting cognitive function with high demand of neural resources in normal aging ^36^. The integrity of frontal regions has also been proposed as pivotal feature for successful cognitive aging ^37^. In the current study, the association between the gray matter volume of right FP and VSTM was primarily driven by individuals with a larger gray matter volume of the left MTL within the normal control group. This reflects better allocation of neural resources and integration between the frontal area and MTL to support cognitive function in normal elderly individuals who have better-preserved brain structure.

On the contrary, in individuals with neuropathology, the engagement of frontal area may be associated with a compensatory mechanism aiming at maintaining cognitive function. Previous research has proposed a compensatory role of the prefrontal area in helping to sustain cognitive function in aMCI when hippocampus-related function is impaired ^38,39^. It has been suggested that as long as brain damage is not significant enough to hinder the recruitment of additional neural resources, individuals with aMCI can still accomplish cognitive tasks by involving frontal neural resources ^9,40^. This compensatory mechanism has been suggested to be closely correlated with disease severity, following a non-linearly U-shape trajectory ^41-44^.

In the current study, the association between gray matter volume of right FP and VSTM in the aMCI group was mainly driven by individuals with smaller gray matter volume in the left MTL. When we used the atrophic left MTL as a seed, an FP area overlapping with the structurally atrophic right FP exhibited stronger functional connectivity with the exact atrophic left MTL in the aMCI group compared to the NC group. All these findings reflect compensatory neuroplasticity, in which the frontal area may act as a cognitive buffer to compensate for the cognitive decline resulting from gradually worsened MTL dysfunction. The stronger connectivity between the left MTL and the right FP may represent a reconstructed neural mechanism due to MTL dysfunction. As neural degeneration progresses, the VSTM underlying mechanism may rely less on traditionally engaged neural resources. However, it’s important to note that this compensatory mechanism may eventually break down as the disease progresses and more severe structural atrophy occurs. The varying brain-behavior associations observed among aMCI subgroups in the current study provide valuable insights into the role of MTL dysfunction and prefrontal compensatory mechanisms in VSTM along the trajectory of disease pathology. These findings could be a pivotal feature for the development of personalized early diagnosis and intervention strategies for AD pathology ^45^.

There are some limitations in the current study. Firstly, it’s important to exercise caution when interpreting structural and resting-state functional data and when discussing the neural correlates of cognitive performance. The inclusion of task-based fMRI data could provide us a more direct and in-depth understanding of the impact of neural deterioration on VSTM in the preclinical stage of AD. Secondly, the aMCI participants and the NC participants in the current study differed in terms of education duration, gender distribution, and were collected from different sites. While it was challenging to recruit participants with perfect matching across all these variables, we have included them as covariates to regress out their potential influences. However, it’s important to acknowledge that these differences may still have the potential to limit the power and generalizability of our results. Thirdly, our results provided valuable insights into the alteration of brain-behavior associations as the disease progresses. However, it also highlights the need for longitudinal research in this area. Future longitudinal studies with carefully selected samples are expected to provide a deeper understanding of the relationship between the brain function and VSTM, and to explore the possibility of utilizing such brain-behavior associations as indicators for early detection of AD.

## Conclusion

The present study highlighted the “MTL dysfunction, frontal compensation” mechanism in response to the VSTM among individuals with aMCI. Interestingly, even as the left MTL displayed atrophy and could no longer fully support VSTM in aMCI, the atrophic right FP area still maintained an association with VSTM performance. Additionally, the heightened connectivity between the left MTL and right FP reflects neural reorganization in response to brain deterioration by the disease’s pathology. Furthermore, the association between the right FP and VSTM was influenced by the gray matter volume of left MTL, indicating the impact of the impact of disease’s progression trajectory on the compensatory mechanism. Taken together, these findings significantly enhanced our understanding of the pathology during the aMCI stage and provide valuable insights into potential intervention targets for early AD.

## Method

### Diagnostic assessment

in the current study, our participant cohort was recruited from hospital. The diagnosis of aMCI was carried out by neurologists at hospital in accordance with the ‘MCI-criteria for the clinical and cognitive syndrome’ ^46^. This comprehensive diagnostic criteria included the following key elements. (1) Subjective Complaints: Both the participants and their caregivers reported a noticeable decline in memory or cognitive function. (2) Cognitive Assessment: Participants underwent cognitive assessment using standardized tests, specifically the Mini-Mental State Examination (MMSE) or the Montreal Cognitive Assessment (MoCA). Scores were adjusted based on the individual’s level of education. (3) Clinical Dementia Rating Scale (CDR): A CDR score of 0.5 was used as an indicator of MCI. (4) Cognitive Deficits: Participants had to meet at least one of the following three criteria: (1) Performance below established cutoffs (> 1 standard deviation) in at least two cognitive domains; (2) Impairment (> 1 standard deviation) in at least two cognitive domains; (3) Experiencing functional limitations in more than one area of instrumental activities of daily living (IADL-14), with a score of 1 or higher.

These rigorous diagnostic criteria ensured the accurate identification of individuals with aMCI and allowed us to focus our research on a well-defined cohort that represents the transitional stage between normal aging and Alzheimer’s Disease. The application of these criteria in our study strengthens the clinical significance and reliability of our findings.

### Participants

The demographic data for the study participants is summarized in Table 1. A total of 123 participants were recruited from hospital, comprising 55 participants diagnosed with aMCI and 68 age-matched normal elderly controls (NC). The participants’ ages ranged from 60 to 90 years old, and the two groups were carefully matched in terms of age. However, there were noticeable differences in terms of education levels and gender distribution between the two groups, which would be regressed out in our analyses. Importantly, significant differences in MMSE and MoCA scores were observed between the two groups, further highlighting the cognitive distinctions between aMCI and NC participants. Informed consent was obtained from all participants and the study was approved to ensure that the research adhered to ethical and safety guidelines consistent with the principles outlined in the Declaration of Helsinki.

### VSTM task

VSTM was assessed using a modified color change-detection task (Fig. 1) ^27^. An arrow was presented on the top of the black fixation across for 200ms and then disappeared. Then after a 300ms break, a brief bilateral array of colored squares was presented for 500ms and participants were asked to remember the items in only one hemifield, which was indicated with the arrow. After a 900ms retention interval, memory was tested with the presentation of a test array. Participants pressed one of the two buttons to indicated whether the test array was either identical to the memory array or differed by one color. The presentation of test array would last until participants gave their response. Accuracy (ACC) and mean reaction time (RT) were calculated for each participants, and capacity was represented by Cowan’s K coefficient (load x (hit rate -false alarm rate)) ^47^. Repeated measure ANOVA was used to test the between-group difference and significant threshold was set at *p* < 0.05.

### MRI Acquisition

Image data were collected from two sites: sample from site 1 included 76 participants (35 aMCI participants and 41 NC participants) and sample from site 2 included 47 participants (20 aMCI participants and 27 NC participants). There is no significant between-group difference in the site distribution (χ^2^ = 0.144, *p* = 0.704). To delineate the influence of machine, collecting site would be use as covariate in imaging analysis.

Imaging data from site 1 was collected with the following parameters: for structural imaging acquisition, TE = 2.98ms, TR = 2530ms, FOV = 256mm x 256mm, acquisition matrix = 256 x 256, reconstruction voxel size = 0.5 x 0.5 x 1.0 mm^3^. For resting-state functional imaging acquisition, TR = 500ms, TE = 30ms, FOV = 224mm x 224mm, matrix = 64 x 64, 25 interleaved slices, and 960 scans collected.

Imaging data from site 2 was collected with the following parameters: for structural imaging acquisition, TE = 2.98 ms, TR = 2530 ms, FOV = 256mm x 256mm, acquisition matrix = 256 x 256, reconstruction voxel size = 1.0 x 1.0 x 1.0mm^3^. For resting-state functional imaging acquisition, TR = 2000ms, TE = 30ms, FOV = 224mm x 224mm, matrix = 64 x 64, 31 interleaved slices, and 240 scans collected.

### Voxel-based morphometry (VBM) analysis

T1-weighted images were manually aligned to conventional anterior commissure (AC) – posterior commissure (PC) space with landmarks including the AC, PC and midsagittal plane and analyzed with CAT toolbox implemented (http://www.neuro.uni-jena.de/cat/) in Statistical Parametric Mapping (SPM12) software (https://www.fil.ion.ucl.ac.uk/spm/software/spm12/). The analysis procedure included skull stripping, segmentation of images into gray matter, white matter and cerebrospinal fluid probability images, and spatial normalization of the gray matter images to a customized gray matter template in standard Montreal Neurological Institute (MNI) space. Gray matter probability maps were held at the threshold of 0.1 to minimize inclusion of incorrect tissue types. The images were modulated with Jacobian determinants and after segmentation they were smoothed with an isotropic Gaussian Kernel (8mm full-width half maximum [FWHM]).

Two-sample t-test was performed to test the between-group difference with total intracranial volume (TIV), age, gender, education and collecting site as covariates of no interest. Statistical threshold was set as height threshold of *p* < 0.005, extent threshold of *p* < 0.05 by using Gaussian random field theory (GRF) in DPABI toolbox ^48^. Significant clusters were determined after applying gray matter mask of AAL (Anatomical Automatic Labelling) 90 regions ^49^. Anatomical locations of brain regions were identified by AAL ^49^, Harvard-Oxford atlas ^50^, and Juelich histological atlas ^51^. Follow-up Pearson correlation analysis examined the relationship between gray matter volume of regions of interest (ROIs; clusters identified from whole brain analysis) and VSTM, controlling for the TIV, age, gender, education and collecting site.

### fMRI data preprocessing and connectivity analysis

The functional images of first ten seconds were discarded to allow for signal equilibrium. Images were preprocessed and analyzed using CONN toolbox ^52^. The preprocessing procedure included following steps: realign and unwrap, slice timing, segmentation and normalization, and 8mm FWHM Gaussian kernel smoothing. 9 participants in aMCI group and 4 participants in NC group were excluded due to excess head motion (displacement > 3mm or rotation > 3°). As a result, 46 aMCI participants and 64 NC participants were included in the fMRI analysis. In addition to the six motion parameters and their first derivatives, WM signals and CSF signals were removed with CompCor method ^53^. This component-based noise correction method reduces physiological and extraneous noise and provides interpretative information on correlated and anticorrelated functional brain networks. Detrend and a 0.008-0.08 Hz band-pass filter were also applied.

In the functional analysis, we would focus on the brain regions showing association with VSTM in the structural analysis. Such areas were used as seed regions of interest (ROI seeds) in the functional connectivity comparison analysis. Seed-to-voxel functional connectivity analysis was performed to identify the functional connectivity of the ROIs with the rest of the brain. Two-sample t-test was then performed using CONN toolbox to investigate whether functional circuits of the ROIs were different between the two groups, with age, gender, education and collecting site as a covariate of no interest. Statistic threshold was set at height threshold of p < 0.005 and extent threshold of p < 0.05 by Gaussian random field theory (GRF) in the DPABI software ^48^. Significant clusters were determined after applying gray matter mask of AAL (Anatomical Automatic Labelling) 90 regions ^49^. Anatomical locations of brain regions were identified by AAL, Harvard-Oxford atlas, and Juelich histological atlas. Follow-up Pearson correlation examined the relationship between the difference of functional connectivity circuits of the ROIs and VSTM, controlling for age, gender, education and collecting site.

### Exploring the interaction between MTL and frontal area

In each group, participants were split into two subgroups, larger gray matter volume of left MTL (LV subgroup) and smaller gray matter volume of left MTL (SV subgroup) by comparing with the median of gray matter volume of left MTL. The correlation between VSTM performance and the gray matter volume of right FP was conducted with Pearson correlation analysis in each subgroup. TIV, age, gender, education and collecting site were used as covariates.

