## Supplementary Figure 1 for "Neural Deterioration and Compensation in Visual Short-term Memory Among Individuals with Amnestic Mild Cognitive Impairment"

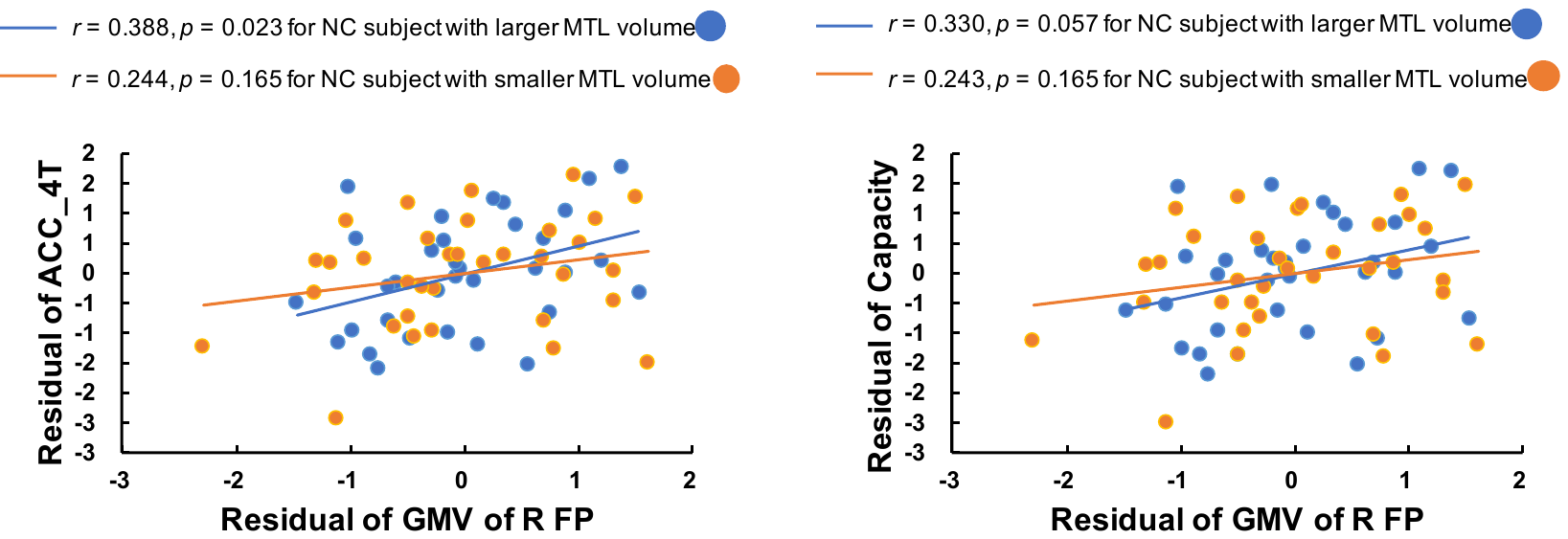

**Supplementary Figure 1.** Relationship between gray matter volume (GMV) of right FP and VSWM performance influenced by the GMV of left MTL in the NC group. Abbreviation: NC, normal control; FP, frontal pole; MTL, medial temporal lobe; L, left; R, right.
